# Differentiating wild and domesticated enset (Musaceae) using phytolith analysis

**DOI:** 10.1101/2024.09.09.611979

**Authors:** Cristina Cobo Castillo, Alemseged Beldados, Philippa Ryan, Sandra Bond, Luc Vrydaghs, Ermias Lulekal, James Borrell, Harriet Hunt, Dorian Q Fuller

**Affiliations:** University College London, Institute of Archaeology, U.K; Department of Anthropology, University of North Carolina, Wilmington (UNCW), USA; Royal Botanic Gardens, Kew, U.K; Vrije Universiteit Brussel, Belgium; Addis Ababa University, Ethiopia

**Keywords:** archaeobotany, Ensete, Ethiopia, Musa, phytoliths, plant domestication, vegeculture, volcaniforms

## Abstract

Enset (*Ensete ventricosum,* Musaceae) is an important economic crop from Ethiopia which accounts for 20% of the staple diet in Ethiopia today. However, its evolutionary history and spread is poorly understood. Archaeology could provide evidence of past use and contribute to our understanding of its early history, but so far, this has not transpired. Cultivated enset is clonally reproduced and seed production rarely occurs, therefore, looking for seed remains is futile and instead archaeobotanical research should focus on microfossils such as phytoliths. Phytoliths have been shown to be diagnostic for the presence of banana (*Musa*) and are expected to be similarly useful for identifying enset, but we need a better understanding of phytolith production and variability, and the extent to which this may be used to track domestication. The current study provides a fundamental baseline for the identification of *Ensete* phytoliths through the examination of phytoliths from leaves and other plant parts based on their size and shape. We consider the differentiation of phytoliths across a single plant, based on location in the leaf, the age of the leaf, and different organs of the plant. We also compare phytoliths in the Musaceae Family, and between the enset cultivar and wild samples.

**HIGHLIGHTS:** We provide the first identification baseline to differentiate wild from domesticated enset by looking at variation across an individual enset plant and comparing these results to phytoliths from wild plants.

## INTRODUCTION

The importance of *Ensete ventricosum* (Welw.) Cheesman (enset) as a food and non-food resource in Ethiopia has long been acknowledged, but its evolutionary trajectory, history and spread is not well understood (Finneran 2007). This paper sets out criteria to determine the presence of enset in archaeological contexts to trace the evolution and spread of this important African staple crop. We provide criteria to identify domesticated and wild enset in archaeological samples by examining their phytoliths, plant microremains in the form of silica bodies found inside plant tissues. Phytoliths are useful for archaeologists because they have excellent preservation and recovery potential in almost all environments. When the plant dies the inorganic silica remains. If plant parts are burned, phytoliths remain as a component of ash deposit.

Enset is a large perennial herbaceous plant belonging to the Musaceae family. Mature plants grow to approximately 6-12 m height, with a wide swollen pseudostem that can reach 1 m in diameter and large paddle-shaped leaves that grow in a spiral (Birmeta et al. 2004). Enset is monocarpic and dies after fruiting, although enset cultivated for food is rarely allowed to fruit and instead is harvested before flowering. In the wild, it is pollinated by bats and the primary seed dispersal mechanism is by rodents and primates (Bekele-Tesemma et al. 1993; Birmeta et al. 2004; Ssali and Sheil 2019). Enset is also referred to as fibre banana, wild banana, Abyssinian banana, Ethiopian banana, or false banana in English (Kelecha 1987). These names make reference to its close relative, the banana (*Musa*), which is similar in appearance. Like the banana, enset has multiple uses besides food which include fodder, medicine, serving plates, a cover while baking bread, wrapping material, and raw material for crafts and construction. However, these two domesticates have distinct culinary uses and processing stages. Enset’s most important role is as a starchy staple food (Murdock 1960), and whereas the primary edible part of the banana is the fruit, enset fruit is not eaten and instead, the corm and pseudostem are consumed.

Although banana and enset are morphologically similar, one produces one of the world’s most popular fruit whilst the other is a virtually unknown crop outside of Ethiopia. Bananas are consumed globally and in 2022, total worldwide export quantity was 19.3 million tonnes (FAO, 2024). On the other hand, enset cultivation and consumption is concentrated in south and southwestern Ethiopia. Enset is the staple food for 20% of the Ethiopian population and feeds ∼20 million people per year (Borrell et al. 2019; Zeberga et al. 2014). It is considered a food security crop due to its drought tolerant nature and high productivity (Brandt et al. 1997; Zeberga et al. 2014). A smaller land area of enset cultivation is found in central Ethiopia, but in the 14^th^ to early 20^th^ centuries CE, there were more areas of cultivation in the north (Bruce 1790; Stiehler 1948; Simoons 1960, Pankhurst 1961). The first mention of enset in northern Ethiopia was in the 14^th^ century CE by the Egyptian writer Ibn Fadl Allah. In the 16^th^ century CE, the Portuguese missionary Manuel de Almeida observed and documented it growing at *Azazo*, Begemeder (present day Gondar). Wild enset populations range from north to south Ethiopia (Fig. 1), as well as elsewhere in Africa (Borrell et al. 2019).

**Fig. 1.**
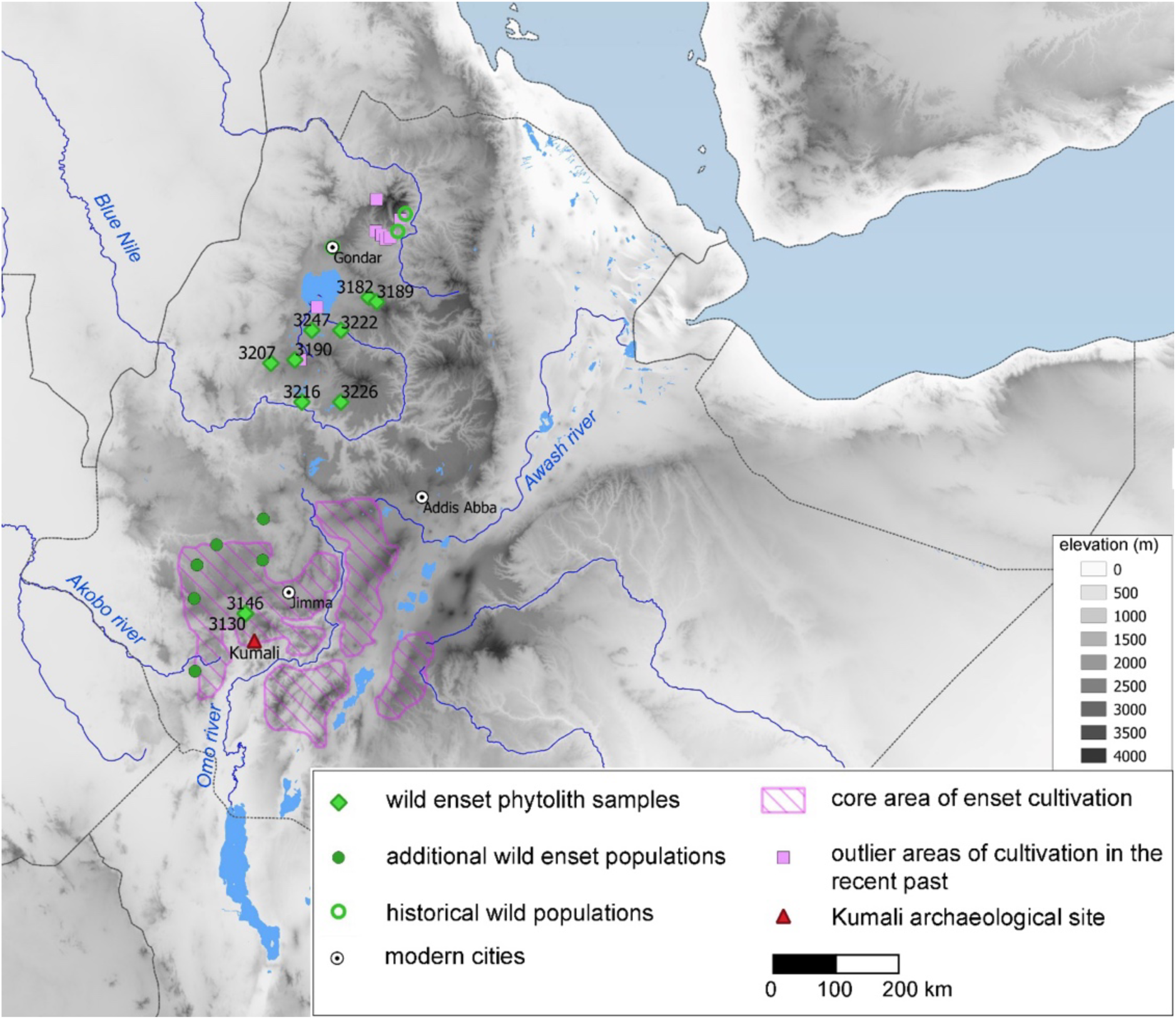
Map of sampled populations of wild enset sampled in this study (Table S1) in relation to the broader distribution of wild enset in Ethiopia (based on White et al. 2023; Brandt et al. 1997; Simoons 1960), the core area of enset cultivation (after Borrell et al. 2019) and outlier areas of cultivation in the recent past (Simoons 1960; Brandt et al. 1997). Also shown is the location of the archaeological site of Kumali with ancient Musaceae leaves (Hildebrand et al. 2010).

Domesticated enset originated in the Ethiopian Highlands, an area regarded as a centre of crop diversity and domestication (Harlan 1969; Harrower et al. 2010; Fuller and Hildebrand 2013; Beldados et al. 2023). Its high levels of intra-crop diversity have been attributed to environmental factors, such as elevation, and human interaction (Samberg et al. 2010). Recent genomic modelling of divergence between wild and domesticated enset suggests that this took place in a single region and cultivation began with sexually reproducing populations rather than resulting from clonal propagation from selected wild plants (White et al. 2023). This implies that much of the recent distribution of enset cultivation has resulted from the dispersal of crops from an original centre of domestication. When and where this dispersal process began and proceeded is an unresolved problem for archaeological research. Archaeological evidence for enset is poor and so far, the only evidence for its antiquity is the desiccated Musaceae leaves from Kumali Rockshelter reported from layers above a radiocarbon date that falls between 250-335 CE (Hildebrand et al. 2010). These leaves were found together with coffee seeds but have not been analysed further to attribute them to the *Ensete* genus (Hildebrand et al. 2010). Since clonally cultivated enset is rarely expected to produce seeds, the routine study of archaeological seed assemblages is unlikely to contribute to the early history of cultivation. Instead, the most feasible approach to documenting past use of enset is through phytolith analysis from archaeological sites. Phytoliths have been shown to be diagnostic for the presence of *Musa* (Mbida et al. 2001; Ball et al. 2006; Lentfer 2009; Vrydaghs et al 2009) and are expected to be similarly useful for identifying *Ensete*. Thus, to progress the archaeobotanical recovery of enset, we need a better understanding of phytolith production and variability, and the extent to which this may be used to track domestication. The current study provides a fundamental baseline for the identification of *Ensete* phytoliths.

Phytolith studies focused on African crops including enset are sparse. Work has focused on millets (e.g. Out and Madella 2016; 2017) and wild grasses (e.g. Fahmy 2008; Neumann et al. 2017; Le Moyne et al. 2023). Large silica accumulators in the monocotyledons belong to the Poaceae, Cyperaceae, and Arecaceae families and the Zingiberales order that includes Musaceae (Prychid et al. 2003; Ge et al. 2022). In this paper, we provide results on phytolith production and variability to refine phytolith identification of enset vis-à-vis wild enset, and other Musaceae. We also investigate whether the different parts of the enset plant, as well as the processed food products contain phytoliths and if these morphotypes are diagnostic. To do this, we first studied the morphology and morphometry of the phytoliths from a single domesticated enset plant, then those from leaves of several wild enset and other species from across the Musaceae family. Morphological and morphometric variation of phytoliths were noted and these provided a baseline for a working description of the phytoliths found in *Ensete* spp., *Musa* spp. and *Musella* sp. The results are intended to define identification criteria to help archaeobotanists analysing sediments from archaeological sites where enset may be present, such as at Kumali rockshelter or Mota and Tuwatey caves (Hildebrand et al. 2010; Ruiz-Giralt and Beldados 2024), and to identify situations when enset microfossils will most likely preserve. This will allow for a better understanding of the historical uses of this crop. We successfully demonstrate that the sampled domesticated enset reliably segregates from wild enset and other Musaceae species, and these can be identified as such.

We propose that one can identify domesticated from wild enset through the examination of leaf phytoliths based on their size and shape. Furthermore, domesticated enset leaf phytoliths differ from *Musa*, as originally suggested in Perrier et al. (2011). Morphotypes or phytolith variants in *Musa* have already been noted in previous studies, including semi-quantification of the frequency of different variants in each species (Ball et al. 2006; Vrydaghs et al. 2009). According to Vrydaghs et al. (2009), there are at least five volcaniform variants per species sampled. Chen and Smith (2013) also show the variability of phytoliths in the order Zingiberales, including three morphotypes in the Musaceae family depending on whether these are from vegetative or reproductive organs. The present study, however, considers in greater detail the differentiation of phytoliths across a single plant, based on location in the leaf, the age of the leaf, and different organs of the plant.

## MATERIALS AND METHODS

### Sampling

Wild enset leaves were collected during fieldwork in Ethiopia in 2020 as part of genetic research (White et al. 2023; Fig. 1). The samples were placed in envelopes with dry silica and transferred to Kew Gardens for sequencing. Leaves from ten wild enset plants were subsampled for phytolith extraction, covering a range of geographic distribution (Fig. 1). Elevation of these collections ranged from 1680 to 2720 masl, extending the elevation ranges from previous publications (Bekele-Tesemma et al. 1993; Eshetae et al. 2021; Friis 1992). These cover the upper part of the wild enset range, which is generally reported to occur between 1300 and 2300 masl (Borrell et al. 2019). This range overlaps with that of cultivated enset (mostly 1500-3000 masl). Age and size of the ten plants also varied. Although not recorded for every accession, wild enset heights range from 220-800 cm, with basal circumferences from 67-250 cm. The largest specimen (height 800 cm, circumference 250 cm) was reported to be ∼6 years old. For those enset plants with size measurements (n=5) there is a clear correlation between height and stem size, which allows some approximate estimation of age (Table S1). Age of most wild enset was not known but was inferred to range from 2 to 4 years for four specimens. A map and list of samples is found in Fig. 1 and Table S1. For the domesticated enset, a 5-year-old flowering plant from the Temperate House at Kew Gardens was sampled (Table S2). We sub-sampled as many plant parts as possible from this one individual, including the leaves, pseudostem, corm, infructescence, bracts and roots. From the youngest leaf, ten leaf cuttings were taken of which seven were processed for phytoliths. There were 57 leaf cuttings taken from the oldest leaf and eleven were processed (Table S2). Visual maps of sampled areas of the leaves were drawn (Fig 5A and 5B). Furthermore, the leaves of three other *Ensete* species, six *Musa* species and *Musella lasiocarpa* from the Temperate House and Palm House in the Royal Botanic Gardens, Kew were sampled. A leaf cutting of *Musa sikkimensis* was collected in 2014 from the Jersey ‘Gardens of Samores Manor’. Tables S2 and S3 include the information of plant parts sampled and the measurements.

Sampling of domesticated enset took into account different uses in terms of plant parts and food types. The main use of *E. ventricosum* is as a food source and the main food types are kocho, bulla and amicho. Different parts of the plant are used in the preparation of these food types (Fig. 2). Kocho is a fermented product from the scraped pseudostem (leaf sheaths) and pulverised corm. Bulla is a white powder which comes from the dried liquid of the scraped pseudostem and pulverised corm and is a by-product of kocho preparation. Finally, amicho is the stripped corm of young enset, which is boiled like other starchy vegetative crops, such as taro or potato. The samples of processed enset used for food (kocho, bulla) were taken from the February and May 2023 field seasons. In this study, kocho samples are divided into several main types: 1) scraped pseudostem and grated corm, liquid drained and ready for fermentation; 2) fermented kocho dough; 3) uncooked kocho; and 4) cooked kocho. The two bulla samples were bought from markets at Mizan Teferi and Sodo. Two samples of corm were studied, one from the Temperate House living collection and the other from field collections in Sheko.

**Fig. 2.**
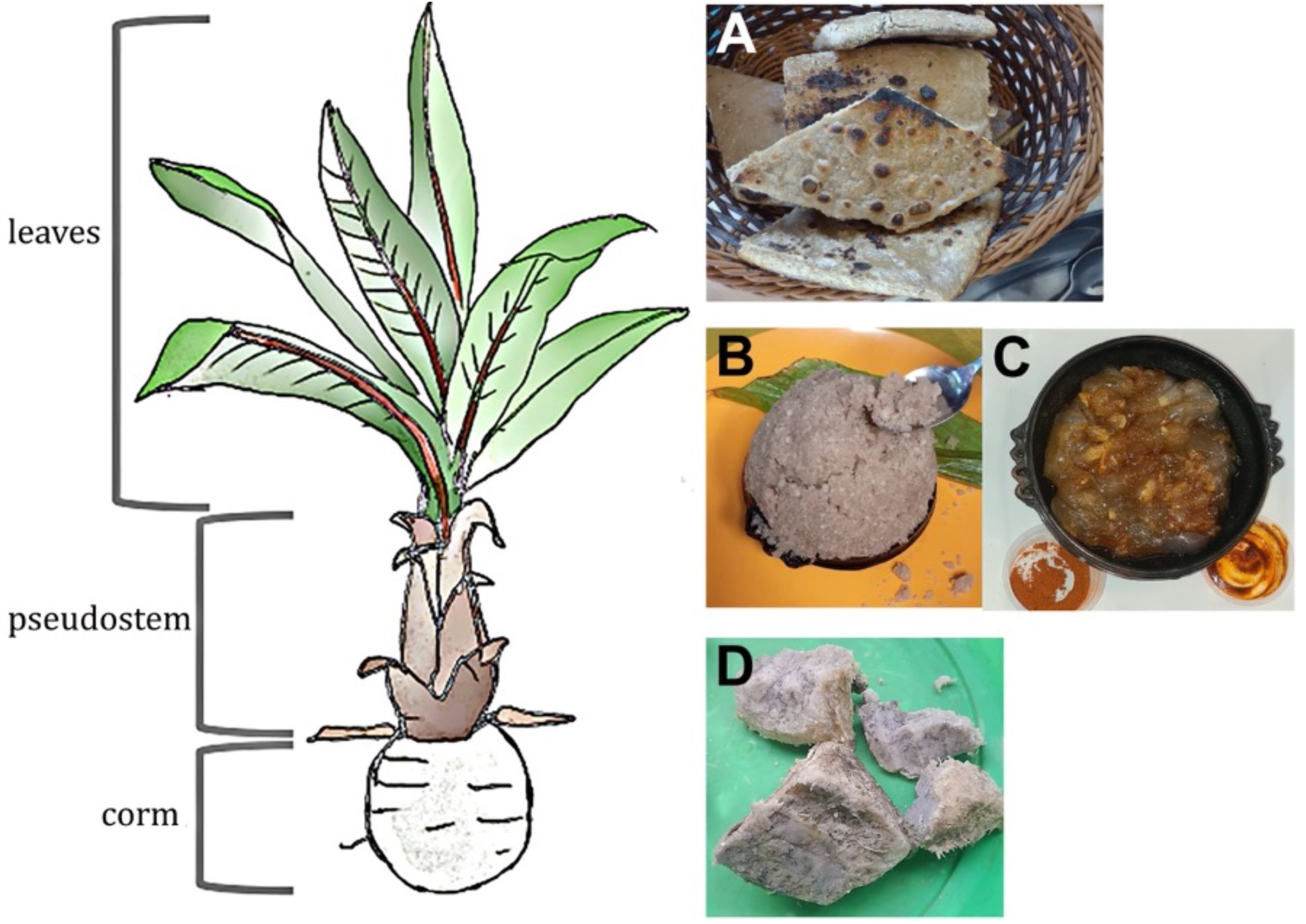
Drawing of enset showing the three main parts of economic value, and at right, examples of the primary enset products. A) kocho, B) bulla cooked as firfir / scrambled from the Sidama Region in south Ethiopia, C) bulla cooked as spiced porridge with pepper and chilli powder and D) amicho from Kambata-Tambaro zone in southern region of Ethiopia (Photos: A - Alemseged Beldados, B - Amesias Alemu, C – Rahel Yitbarek, D) Amesias Alemu).

### Phytolith extraction

Phytoliths from modern plant material were extracted using the dry-ash method (also known as spodograms; Piperno 2006). A ∼2×2 cm section of the plant was cut, washed with deionised water, placed inside a crucible, and dried at 50°C. These were ashed in a muffle furnace at 500°C for 2 hours and allowed to cool inside the furnace. The ash was transferred to tubes and a small amount of 10% HCl was added to eliminate carbonates. Deionised water was added to wash the HCl solution, and the tubes were centrifuged at 2000 rpm for 5 minutes. The solution was decanted with a pipette leaving the phytoliths at the bottom. Deionised water was added, and the tubes centrifuged at 2000 rpm for 2 minutes. This last step was undertaken three times. The solution containing the phytoliths was pipetted onto slides and once dry, mounted with Entellan solution and a cover slip. The same solution containing phytoliths was also mounted onto SEM stubs.

### Imaging and Measurements

The slides were observed under a high-powered transmitted light microscope (Nikon Eclipse LV100). Images and measurements of phytoliths were taken with the x100 oil immersion lens using the GX Capture software. Images which produced finer resolution were acquired using the Hitachi S3400 SEM located in University College London, Institute of Archaeology.

Length and width measurements of approximately fifty phytoliths per sample were taken from articulated phytoliths (see Table S3 for dataset). The area of the phytolith is calculated as length*width and labeled as LxW.

Measurements (LxW) used in the figures (Fig. 6 and 7) correspond to the following and raw data is found in Table S3.
Young leaf base – a total of 55 measurements from sample Temp 67, slide F9.
Young leaf apex – a total of 49 measurements from sample Temp 59, slide F7.
Young leaf all – a total of 363 measurements from samples Temp 59 (slide F7), Temp 60 (slide F17), Temp 62 (slide F13), Temp 63 (slide F10), Temp 64 (slide F14), Temp 65 (slide F15), Temp 67 (slide F9).
Old leaf base – a total of 57 measurements from sample Temp 30, slide F2.
Old leaf apex – a total of 52 measurements from sample Temp 33, slide F5.
Old leaf all – a total of 641 measurements from samples Temp 2 (slide K1), Temp 3 (slide F1), Temp 4 (slide F19), Temp 30 (slide F2), Temp 31 (slide F3), Temp 32 (slide F4), Temp 33 (slide F5).
Wild leaf ALL – a total of 355 measurements from samples #3130, #3190, #3189, #3207, #3146, #3182, #3222.
Musaceae species: *Musella lasiocarpa* (n=47), *Musa balbisiana* (n=49), *M. basjoo* (n=47), *M. beccarii* (n=51), *M. rubra* (n=50), *M. velutina* (n=52), *Ensete glaucum* (n=56), *E. lecongkietti* (n=53), *E. superbum* (n=52), *Musa* ‘ice cream’ ABB (n=52), *E. ventricosum* (n=1004).

### Nomenclature

We use the ICPN 2.0 for phytolith nomenclature to describe the morphotypes. However, we continue to use the *nomen conservandum* ‘volcaniform’ instead of ‘cavate’ in nomenclature of ICPN 2.0 (see Fig. 3) for the key morphotype from Musaceae.

**Fig. 3.**
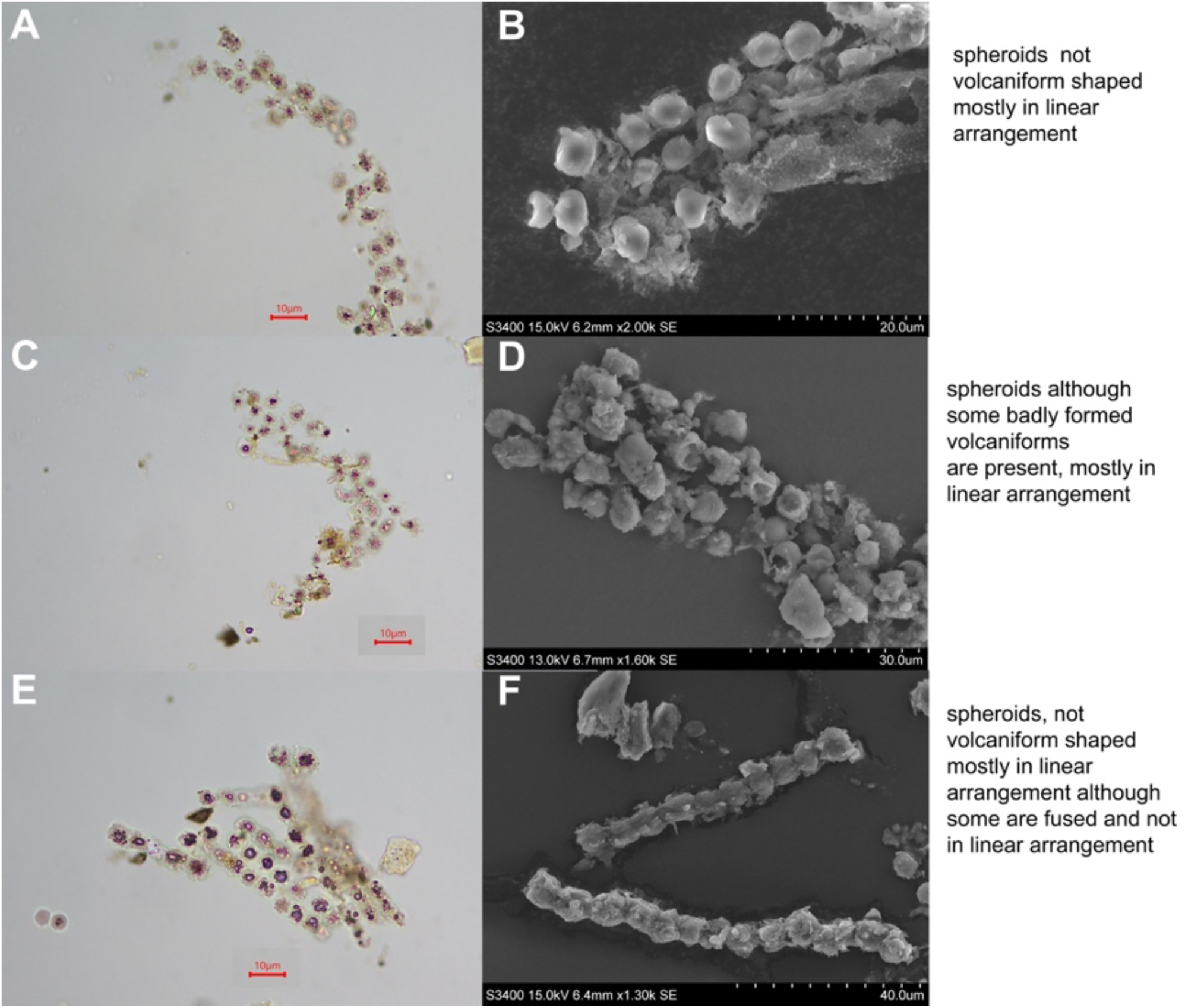
A selection of images of the phytoliths from wild enset and their corresponding general observation. A, C and E images using x100 oil immersion lens (x1000 magnification). B, D and F micrographs using the SEM. A, B - #3130 (estimated age: 3-4 years); C, D - #3190 (age: 6 years); E, F - #3189 (estimated age: 2 years).

## RESULTS

### Wild enset

Leaf samples taken from ten wild enset plants were examined for phytoliths. The phytoliths are SPHEROID ORNATE arranged linearly and do not have the typical volcaniform shape associated with Musaceae except for one sample which exhibited partially formed craters only visible with the SEM at a magnification of x1600 (Fig. 3D). Among those specimens with size/age information this is the largest and oldest wild enset specimen with an informant estimated age of 6 years. Except for one sample belonging to a young wild enset with an estimated age of 2 years (ID#3189; Fig. 3F), none of the other phytoliths from wild enset were joined. No phytoliths were observed in two of the samples.

### Domesticated enset

Phytoliths were identified in the leaves, pseudostem, corm, and seeds (Fig. 4; Table 1) but not in the roots (cf. Tomlinson 1959 for *Musa*). Several morphotypes were observed depending on the part of the plant examined, and the ages of the leaves. The corm and pseudostem phytoliths are SPHEROID ORNATE and BLOCKY respectively, with at least four variants occurring in the pseudostem (Fig. 4C). Seeds produced POLYHEDRAL phytoliths in the testa (Fig. 4F).

**Fig. 4.**
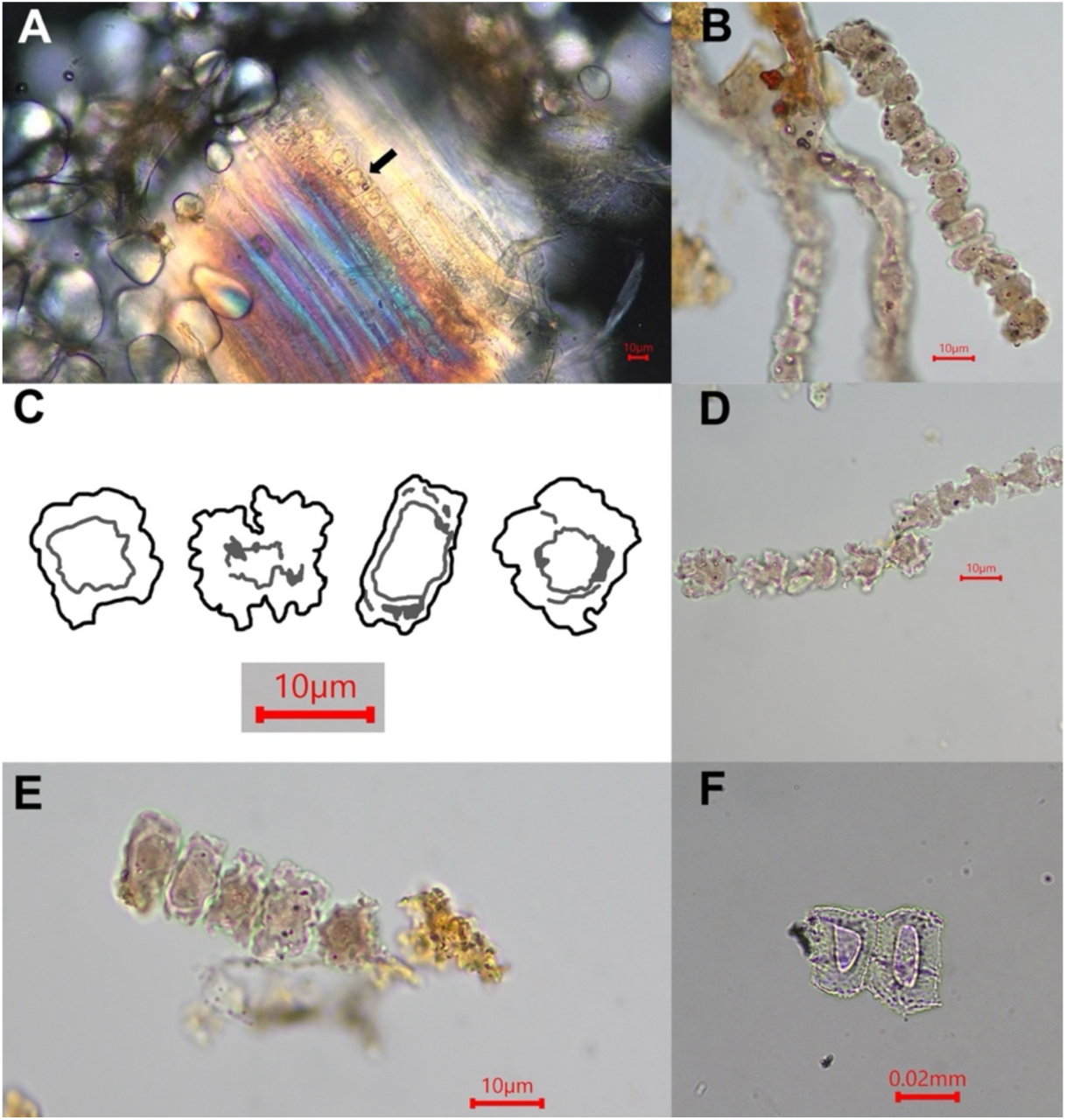
Images of phytoliths from enset plant organs. A) Thin section of corm from Sheko field sample showing a line of SPHEROID ORNATE phytoliths located along a raphide bundle, under polarising filter; starch grains are also visible; B) corm phytoliths from Sheko; C) line drawings of the four BLOCKY morphotypes found in the pseudostem; D-E) pseudostem phytoliths (BLOCKY); F) seedcoat phytoliths (POLYHEDRAL).

**Table 1:**
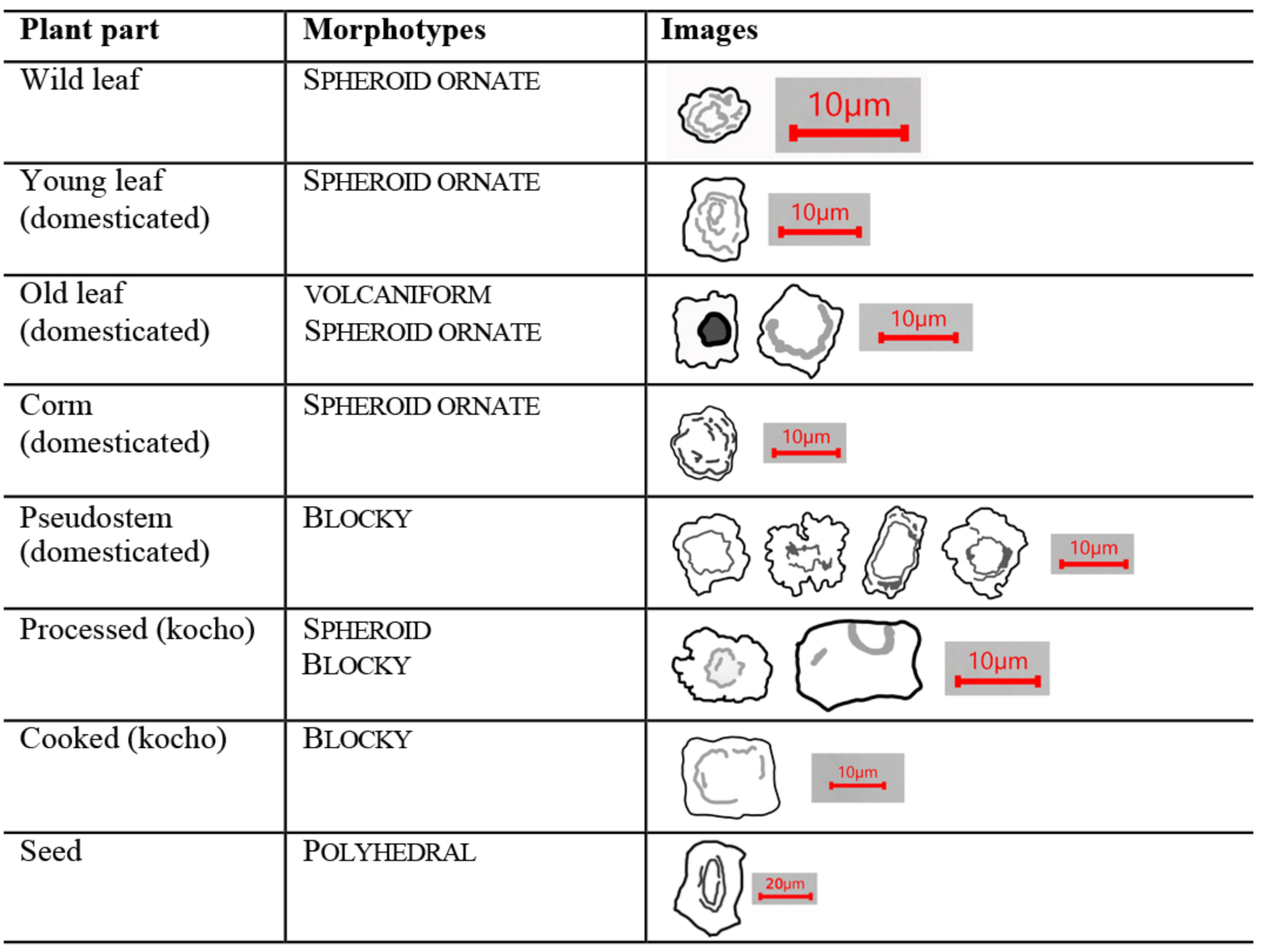
Summary of phytolith morphotypes found in various enset plant parts and processed food products. The images are outlines from the polar view.

| Plant part | Morphotypes | Images |
| --- | --- | --- |
| Wild leaf | SPHEROID ORNATE |  |
| Young leaf (domesticated) | SPHEROID ORNATE |  |
| Old leaf (domesticated) | VOLCANIFORM<br>SPHEROID ORNATE |  |
| Corm (domesticated) | SPHEROID ORNATE |  |
| Pseudostem (domesticated) | BLOCKY |  |
| Processed (kocho) | SPHEROID<br>BLOCKY |  |
| Cooked (kocho) | BLOCKY |  |
| Seed | POLYHEDRAL |  |

The morphotypes in the leaves were dependent on their location and the age of the leaf. The dominant morphotype in the youngest leaf sampled is SPHEROID ORNATE (Fig. 5A) across the entire lamina, like those from wild enset (Fig. 3). Some phytoliths appear as incipiently volcaniform morphotypes, similar to the oldest wild enset. The size of phytoliths found in the base of the young leaf fall within the size range of all wild enset phytoliths and are smaller than those from the older leaf (Fig. 6A). On the other hand, the phytoliths found in the apex of the youngest leaf fall within the size range of the oldest leaf apex (Fig. 6B). Phytolith morphotypes found in the oldest leaf are well-formed volcaniforms except those located at the base and near the base of the leaf, which resemble those found in the youngest leaf and in wild enset, but they are bigger in size (Fig. 6A).

**Fig. 5.**
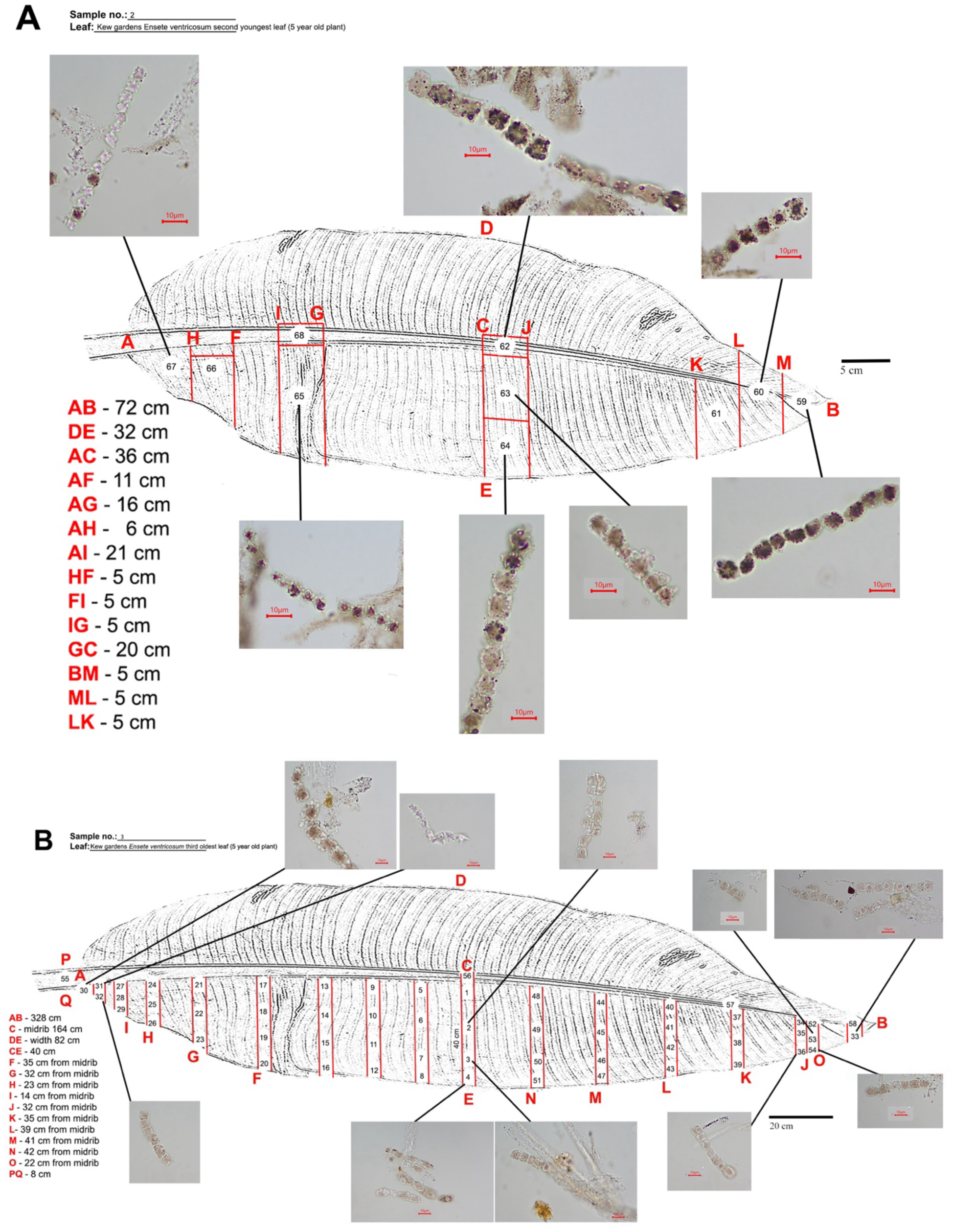
Visual maps showing locations of phytoliths in leaves sampled from the domesticated enset located in the Temperate House, Royal Botanical Gardens, Kew. A: Visual map showing location of phytoliths measured and described in the youngest leaf studied. B: Visual map showing location of phytoliths measured and described in the oldest leaf studied.

**Fig. 6.**
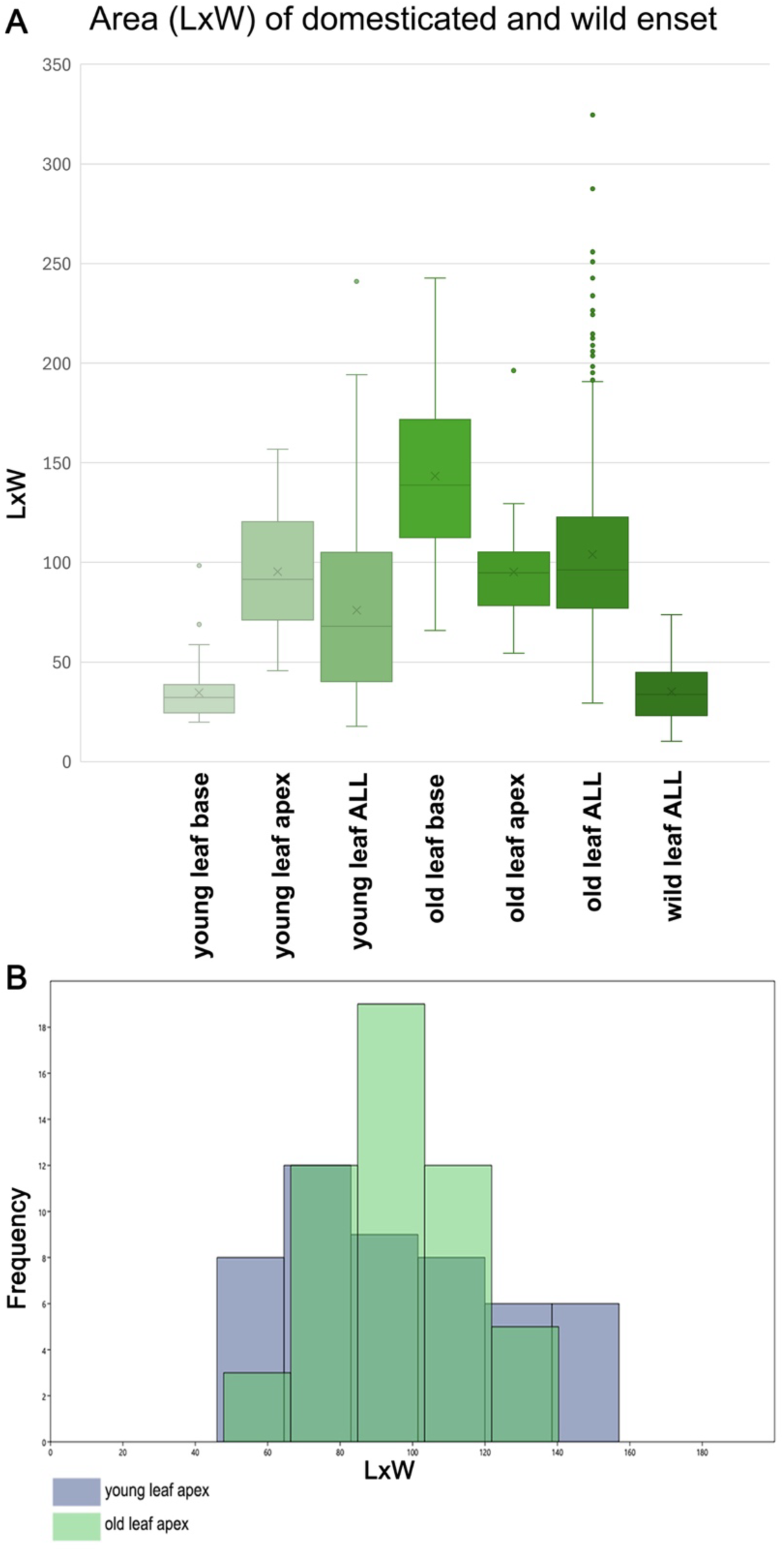
Area (LxW) of leaf phytoliths from domesticated and wild enset. A: Frequency of the area (LxW) of phytoliths from different locations of young and old domesticated enset leaves and those from wild enset. B: Histogram showing the frequency of the area (LxW) of phytoliths from the apex of the young and old leaves of domesticated enset. Young leaf base (n=55), young leaf apex (n=49), young leaf ALL (n=363), old leaf base (n=57), old leaf apex (n=52), old leaf ALL (n=641), wild leaf ALL (n=355).

### Processed enset

Different processing stages of kocho were sampled and showed the presence of phytoliths demonstrating that phytoliths from domesticated enset survive the processing stages of making kocho and are recoverable. The corm and pseudostem produce SPHEROID AND BLOCKY morphotypes and since both these plant parts are used in the production of kocho, the phytoliths found in the processed food should necessarily contain morphotypes from both sources, rather than from the leaf. Kocho contains several phytolith morphotypes identified in the corm and pseudostem: 1) BLOCKY with edges torn off and no craters or, 2) SPHEROID which may be melted or degraded phytoliths. However, they are predominantly BLOCKY. There is a large degree of variation in the sizes and shapes.

The uncooked corm (amicho) from the five-year-old enset in the Temperate House and one from Sheko were examined for phytoliths (Fig. 4A and 4B). The bulla (slide M8) bought from the market had no enset phytoliths. Based on the studied material, preliminary results on processed enset suggest that phytoliths do occur and there is some potential to identify these in enset food preparations, particularly kocho, including from different stages of preparation (drained before fermentation, fermented with fluid residue, after fermentation but uncooked, cooked).

### Comparison of phytoliths from the Musaceae family

Phytoliths from eleven specimens from the Musaceae family, including 3 genera and 11 species, were extracted for morphologic and morphometric comparison (Table 2). The area (LxW) of the phytoliths was plotted and represented in Fig. 7. The results clearly show that the *E. ventricosum* phytoliths are smaller than those belonging to the *Musa* and *Musella* genera. *Musa beccarii*, found only in Borneo, appears to be an exception, although we note that the individual sampled was a very young plant, which could impact phytolith size. In addition to having large phytoliths, the row of leaf phytoliths in *Musa* spp. are more clearly volcaniform than those in wild enset or young leaves of domesticated enset. *Musa* phytoliths are significantly larger than those of *Ensete*, even compared to those from the older *Ensete* leaves. Morphologically, *E. ventricosum* differs from *Musa* spp. in the shape of the protruding cone as defined by Mbida et al. (2001, see Fig. 3). However, phytoliths examined in Mbida et al. (2001) are probably from mature leaves with fully formed phytoliths and no young leaves of *E. ventricosum* domesticates or wild counterparts were examined. Compared to the other *Ensete* genera*, E. superbum* phytoliths are smaller in size than *E. ventricosum* but this species is native to Asia and unlikely to occur in archaeological soil samples in Africa. Two of the Asian ensets, *E. superbum* and *E. glaucum,* produce phytoliths with scalloped edges (like a ravioli) and are morphologically distinct from *E. ventricosum*.

**Fig. 7.**
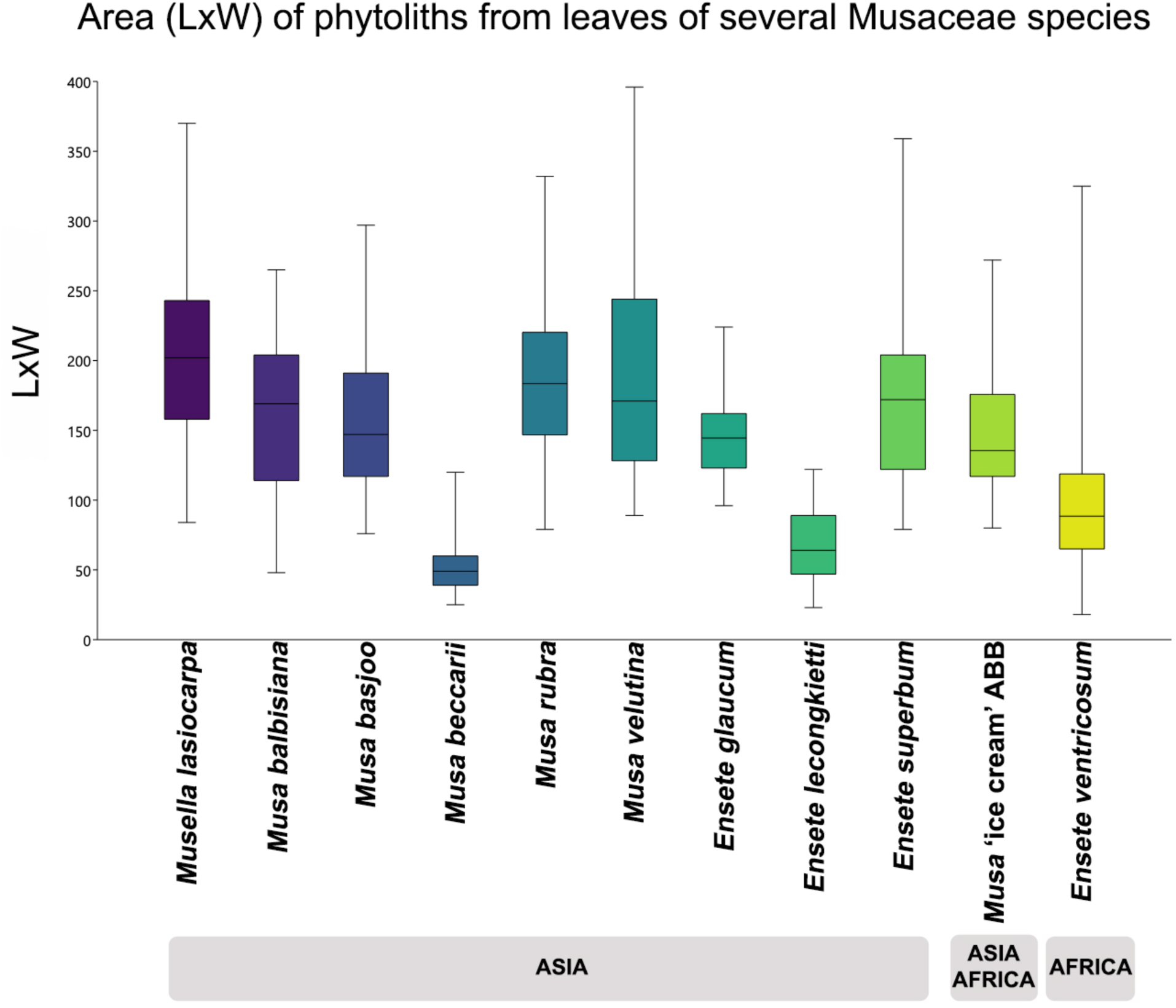
The area (LxW) of leaf phytoliths of eleven species from the Musaceae family, including *Ensete ventricosum*. Information on their geographic origins is included. *Musella lasiocarpa* (n=47), *Musa balbisiana* (n=49), *M. basjoo* (n=47), *M. beccarii* (n=51), *M. rubra* (n=50), *M. velutina* (n=52), *Ensete glaucum* (n=56), *E. lecongkietti* (n=53), *E. superbum* (n=52), *Musa* ‘ice cream’ ABB (n=52), *E. ventricosum* (n=1004).

**Table 2:**
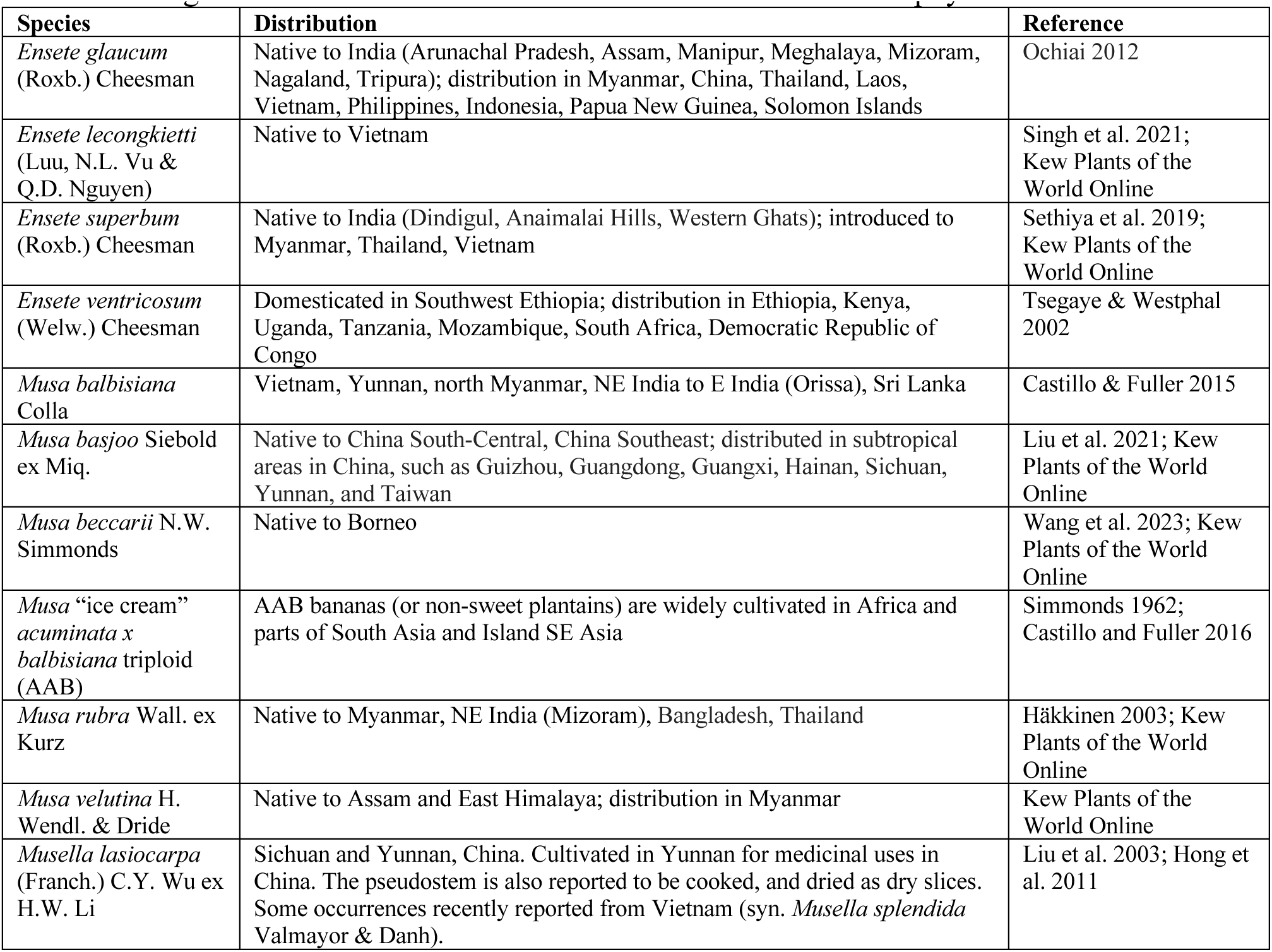
Background information on other Musaceae examined for leaf phytoliths.

## DISCUSSION

Enset phytoliths, as well as *Musa*, are conspicuously underrepresented in African archaeology. Intensively screened samples from sites in the Inner Congo Basin found *Musa* phytoliths in small numbers, whether archaeological or from modern surface samples (Neumann et al. 2022) and none were found from Iron Age sites in southern Cameroon (Eggert et al. 2006). We propose that this may have to do with the locations where the sediment samples are collected but also understanding which plant parts are normally used by people. It has been previously suggested that practices of discarding the phytolith-bearing plant parts, namely the leaves and pseudostems, result in low phytolith counts or limited archaeological signatures since the plant parts are left to decompose in open areas (Neumann et al. 2022; Lentfer 2009). We concur but also suggest that age and the part of the leaf used affects the visibility of phytoliths, as phytoliths in enset are larger, better formed and more silicified in older leaves (and possibly on older plants); similar patterns seem likely for *Musa*.

Although enset is a culturally and economically important crop in Ethiopia, enset use is mostly as a food crop from the corm and pseudo-stem which may leave indeterminate phytolith traces as these are not volcaniform types (see Table 1). Although leaves are useful (e.g., for wrapping food, medicine, craft, traditional mats, and fodder), the younger leaves may be chosen for their flexibility, and any resulting phytoliths from these young leaves are not the classic volcaniform types. The Gurage people from central Ethiopia make a distinction between leaves (*q’t’er*) and the outside layers and dried leaf sheaths (*enewa* and *wedere*) of enset and their specific uses (Hailemariam 1991). *Q’t’er* probably refers to the younger leaves and are used for baking bread, tablecloths, fodder and the different processing stages of enset (Hailermariam 1991). In this study, the researchers observed the use of young leaves for serving food and wrapping kocho. Similarly, young and old banana leaves have different uses. Leaf pruning is a reported practice of enset management in some regions, which is likely to remove leaves before they mature enough to form volcaniforms (Tsegaye and Struik 2002). Thus, while the volcaniforms of older leaves of domesticated enset can be differentiated from wild enset and from *Musa,* they are likely to be rare archaeologically due to patterns of enset use.

A key challenge arises from the indistinct shape or SPHEROID ORNATE phytoliths found throughout the young leaves (Fig. 8). Although phytoliths form in lines above vasculature, as reported for *Musa* volcaniforms (Tomlinson 1959), those found in the young leaves are not volcaniform, but instead are SPHEROID. While Tomlinson (1959) did note the presence of SPHEROIDS in Musaceae, our observation indicates that some of the spheroids in wild enset leaves or young domesticated enset leaves are homologs of *Musa* volcaniforms. While series of joined SPHEROIDS are found in young leaves, these are largely absent from more mature leaves, which instead have chains of discrete well-formed volcaniform phytoliths. We hypothesize that this is because in young leaves the cells are ‘living cells,’ and their cell walls continue to allow osmosis during early phytolith formation. We have successfully demonstrated that phytoliths from young leaves of domesticated enset display smaller sizes, are less volcaniform (less well-formed craters), and less completely silicified with some joined phytolith chains. In our study, the morphotypes of phytoliths from the young leaves of the domesticated enset overlap with wild enset and their sizes and forms (compare Fig. 3, 8 and 6A). By contrast older leaves of the domesticated enset sampled produced diagnostic volcaniform phytolith chains. While the later are diagnostic and can be taken as a mark of domesticated enset, they seem less likely to make their way into the anthropogenic midden deposits which are archaeological features typically sampled for phytoliths. However, more archaeological attention to the presence of SPHEROIDS may potentially signal the presence of enset from archaeological sites, even if these do not separate wild from domesticated forms.

**Fig. 8.**
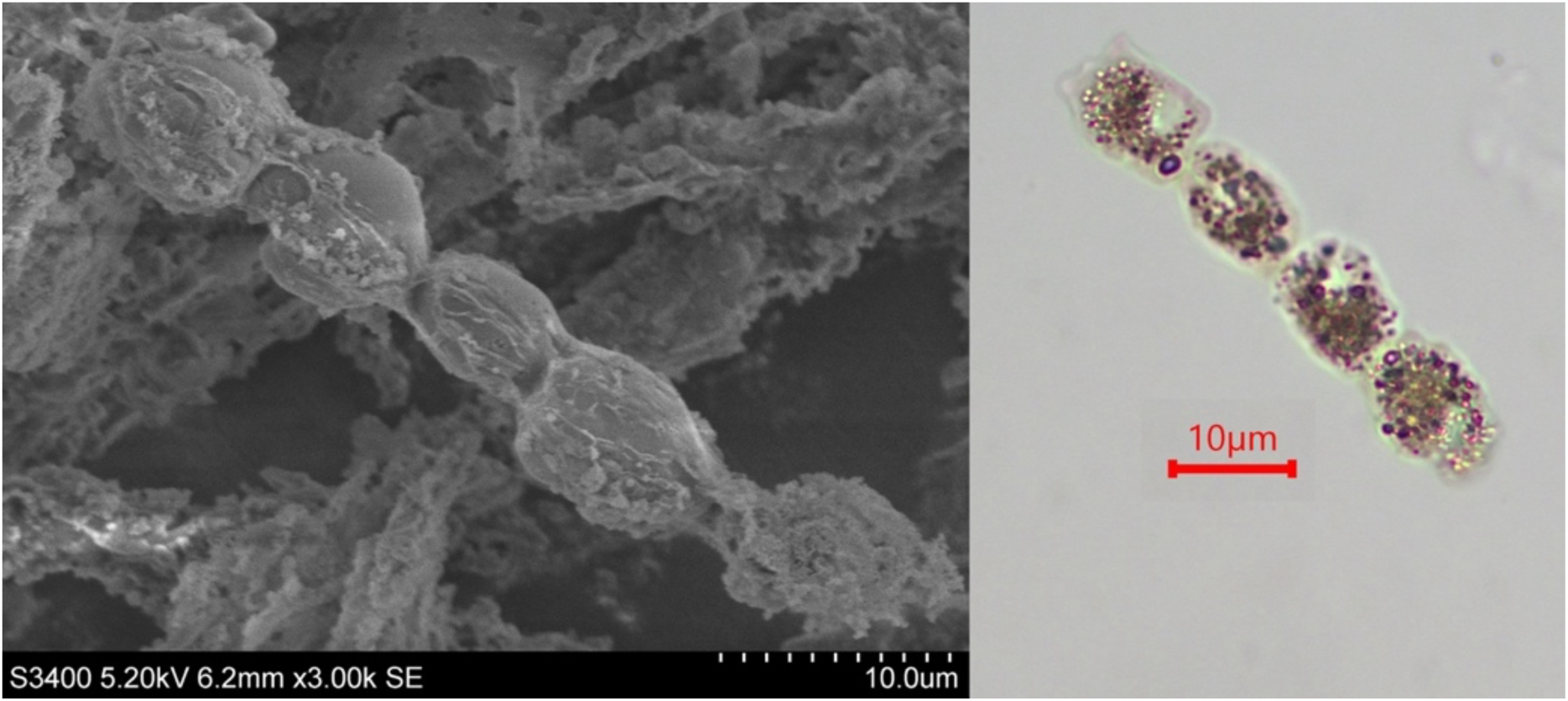
SEM micrograph and high-powered microscope image (x100) of lamina apex phytoliths from the youngest leaf belonging to *Ensete ventricosum* in the Temperate House, Royal Botanical Gardens, Kew. The phytoliths are SPHEROID and joined.

Another subtle distinction between wild and the sampled domesticated enset can be proposed. In the young leaves of the cultivar, the SPHEROID phytoliths are joined (Fig. 8), while they are still undergoing formation in living cells. In contrast, the SPHEROID phytoliths from wild enset were not joined. This probably signifies that the leaves are mature, and the cells are no longer alive. However, processing of archaeological sediments for phytoliths may result in disarticulation, producing phytoliths which appear simply as SPHEROID ORNATE, regardless of whether they came from joined chains or separate cells originally. As has been noted in a previous study, modern sediment samples screened for *Musa* phytoliths yielded very few volcaniform morphotypes associated with Musaceae and are instead dominated by SPHEROID ORNATE, SPHEROID ECHINATE and grass short cell phytoliths (Neumann et al. 2022). It seems a possibility that the SPHEROID ORNATE morphotypes might include those produced by young Musaceae leaves, which calls for further study.

Phytoliths taken from the base of young leaves of domesticated plants overlap in morphometry to those of wild enset phytoliths (Fig. 6A) and are similar in appearance, characterised as SPHEROIDS with uneven edges or SPHEROID ORNATE (Fig. 9A and 9B). As already noted, some/most of the phytoliths from the domesticated young leaf are joined unlike in most wild examples. However, the LW measurements may help to segregate from the wild populations because although the SPHEROID phytoliths of young, domesticated leaves overlap with those of the wild, particularly those from the base of the leaf, they also range to larger sizes than those in wild plants (Fig. 6A). Thus, either of these lines of evidence, joined SPHEROID ORNATE or the presence of well-formed volcaniform would seem to be indicative of domesticated enset. Admittedly, we expect both forms to be much less common than the BLOCKY phytoliths of the edible parts of the plant (Fig. 4).

**Fig. 9.**
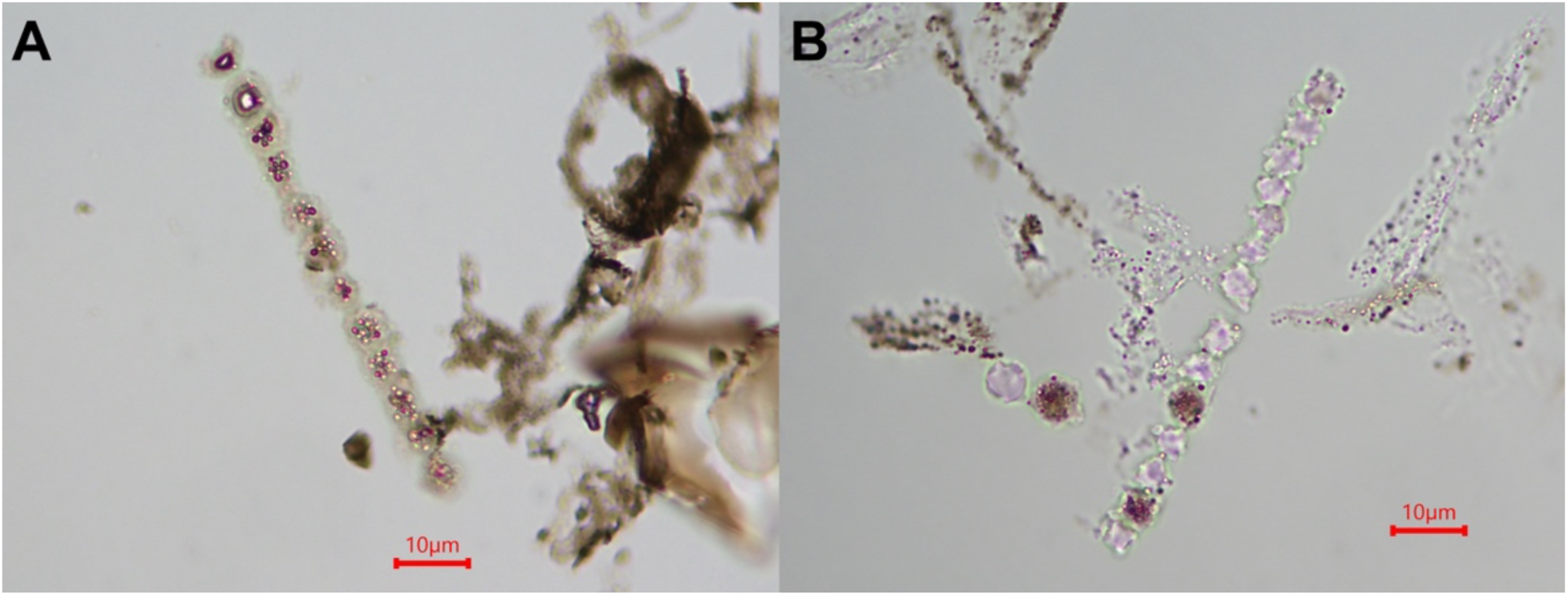
High-powered microscope images (x100) comparing leaf phytoliths from wild and domesticated enset. A: phytoliths from wild enset #3207; B: phytoliths from the base of the youngest leaf studied.

These observations also contribute to understanding some of the ways that domesticated enset evolved away from its wild ancestor. It is already established that enset shared some of the expected domestication syndrome traits of vegetatively propagated crops (Denham et al. 2020), such as an increased yield of edible parts, loss of tuber acridity, and reduced reliance on sexual reproduction. Hildebrand (2003) documents a reduced productivity of seeds in domesticated enset plants allowed to fruit, while her ethnographic informants indicated that corms of wild plants are less palatable. In terms of phytoliths, however, there appears to be a developmental hypermorphy as the derived domesticate has expanded silicification, with better formed volcaniforms reaching larger sizes. Given that wild *E. ventricosum* and *E. lecongkietti* both have small phytoliths compared with taxa across the wider family, we can suggest that smaller phytoliths evolved as a form of paeodomorphosis when these *Ensete* diverged from the ancestral *Musaceae,* because *Musa* and *Musella* share larger, well-developed volcaniforms. The domestication process in *E. ventricosum* then reversed this trend. In archaeological research, the identification of small volcaniform phytoliths within the *E. ventricosum* – *lecongkietti* range, should take into consideration the place of origin and geographical distribution of these plants (Table 2), as well as age of the archaeological sediment studied.

### Conclusion

Originally, leaves from domesticated enset plants were collected by the geneticists at the Royal Botanic Gardens, Kew during fieldwork in Ethiopia in 2020. These specimens were intended to be used for phytolith extraction and examination and to form the basis of the identification criteria for enset. However, the leaves collected for sequencing were usually the younger ones due to the lower content of enzymes compared to older leaves that might affect DNA extraction success. The initial phytolith study using these samples produced inconsistent results, sometimes showing badly formed and shapeless phytoliths resembling those of wild enset. The present study provides an explanation for these results by establishing that phytolith morphology changes due to leaf maturation.

Our work shows how domesticated enset can be identified based on the well-formed, larger volcaniform phytoliths found in older leaves. On the other hand, poorly formed and smaller volcaniforms may be present from younger leaves of cultivars or wild enset plants. Whether wild or domesticated, *E. ventricosum* phytoliths are morphologically and morphometrically distinct from other Musaceae species, especially domesticated *Musa* that were introduced in prehistory from Asia to Africa. It is necessary to use both the shape and size of phytoliths to demonstrate the presence of domesticated enset. Domesticated enset phytoliths, however, are not directly associated with the processing of edible parts, which produce several BLOCKY morphotypes, although these are likely not taxonomically diagnostic on their own. Further work can explore how volcaniforms vary between cultivar groups, altitude, and age of plants. Ethnographic work is needed to assess the assemblage composition and the balance of volcaniforms and BLOCKY morphotypes in the various by-products of human activities. In addition, work across the Musaceae family also requires further work on variability across individual plants, especially in relation to leaf maturity, comparable to the work reported above for *Ensete ventricosum*.

## Supplementary data

The following supplementary data are available at JXB online.

Table S1. Age estimates of wild enset used in this study.

Table S2. Details of plant parts sampled of Musaceae species located in the Temperate House and Palm House, Kew Gardens.

Table S3. Measurements, including length and width, of wild enset, domesticated enset from the Temperate House, Kew Gardens and Musaceae species.

## Acknowledgements

The authors thank David Cooke, Briony Langley and Jessica Francis for their help sampling plants in Kew Gardens, Katharina Neumann for her comments and discussions on phytoliths and enset, Amesias Alemu and Asaye Asfaw for conducting interviews that informed our work, and Jehova Lourenco and Manoj Kumar-Mahto for their help in the laboratory.

## Author contributions

CCC: conceptualization, methodology, analysis, visualization, writing and editing; DQF: writing, review, funding acquisition, and supervision; PR and JB: review and funding acquisition; LV, AB, HH and EL: review and editing; SB: methodology.

